# A novel baseline-effect shift tracking model for more sensitive detection of differences in the effects of closely related dopamine transporter inhibitor/ sigma receptor antagonist drug combinations on psychostimulant use

**DOI:** 10.1101/2024.08.27.609852

**Authors:** Reshma Paul, Roshni M Gandhi, Martin O Job

## Abstract

**Background:** For the purpose of improving the ability to distinguish the activity of closely related drugs on psychostimulant use to enable more specific drug effect characterization, we have developed a new model termed the baseline-effect shift tracking (BEST) model. BEST compares/contrasts the baseline-drug activity relationship(s).

**Aim:** To compare the current model to our BEST model to determine which was more effective in distinguishing the effects of combinations of a dopamine transporter inhibitor (methylphenidate, MPD) and selective sigma1 (BD1063) and non-selective sigma (BD1008) receptor antagonists on cocaine consumption.

**Methods:** Male Sprague Dawley rats were trained to self-administer cocaine (n = 9, 0.32 mg/kg/infusion) or sucrose pellets (n = 6, 20 mg pellets/delivery). We determined the effects for cocaine/sucrose of combinations of MPD (1 mg/kg i.p) and 1) BD1063 (0, 3.2, 10 mg/kg i.p), and 2) BD1008 (0, 3.2, 10 mg/kg i.p) on a) consumption at zero price (Q_0_), and b) essential value (eValue, demand elasticity) estimated using behavioral economic analysis of within-session demand curves, and c) the total intake under the price response curve (TIPR). We compared the models using ANOVA/ regression analysis.

**Results:** The current model did not detect any differences in the effects of these drug combinations on cocaine/ sucrose taking behavior. For cocaine, but not for sucrose, the BEST model detected differences in the effects of these drug combinations on TIPR in subjects with higher baseline activity.

**Conclusion:** BEST model (with TIPR analysis) is more sensitive than the current models in differentiating drug effects on cocaine consumption.

## Introduction

Combinations of dopamine transporter (DAT) inhibitors, such as methylphenidate (MPD), and sigma antagonists, such as BD1008 (a non-selective sigma receptor antagonist) and BD1063 (a selective sigma1 receptor antagonist), are thought to suppress cocaine consumption with similar efficacy (Hiranita et al., 2011; Job and Katz, 2019), for review, see (Katz et al., 2017). However, BD1063 and BD1008 are distinct pharmacologically with regards to sigma receptor selectivity profiles (Figure 1), and as such these drug combinations should have distinct pharmacological effects on cocaine consumption, but this is not clear from current methods. The ability to distinguish the effects of closely related drugs will be an immeasurable asset to the field because it can enable us to develop the best possible drugs to address not only psychostimulant use disorders specifically, but diseases or disorders generally. For this reason, we have developed a new model termed the baseline-effect shift tracking (BEST) model.

**Figure 1.**
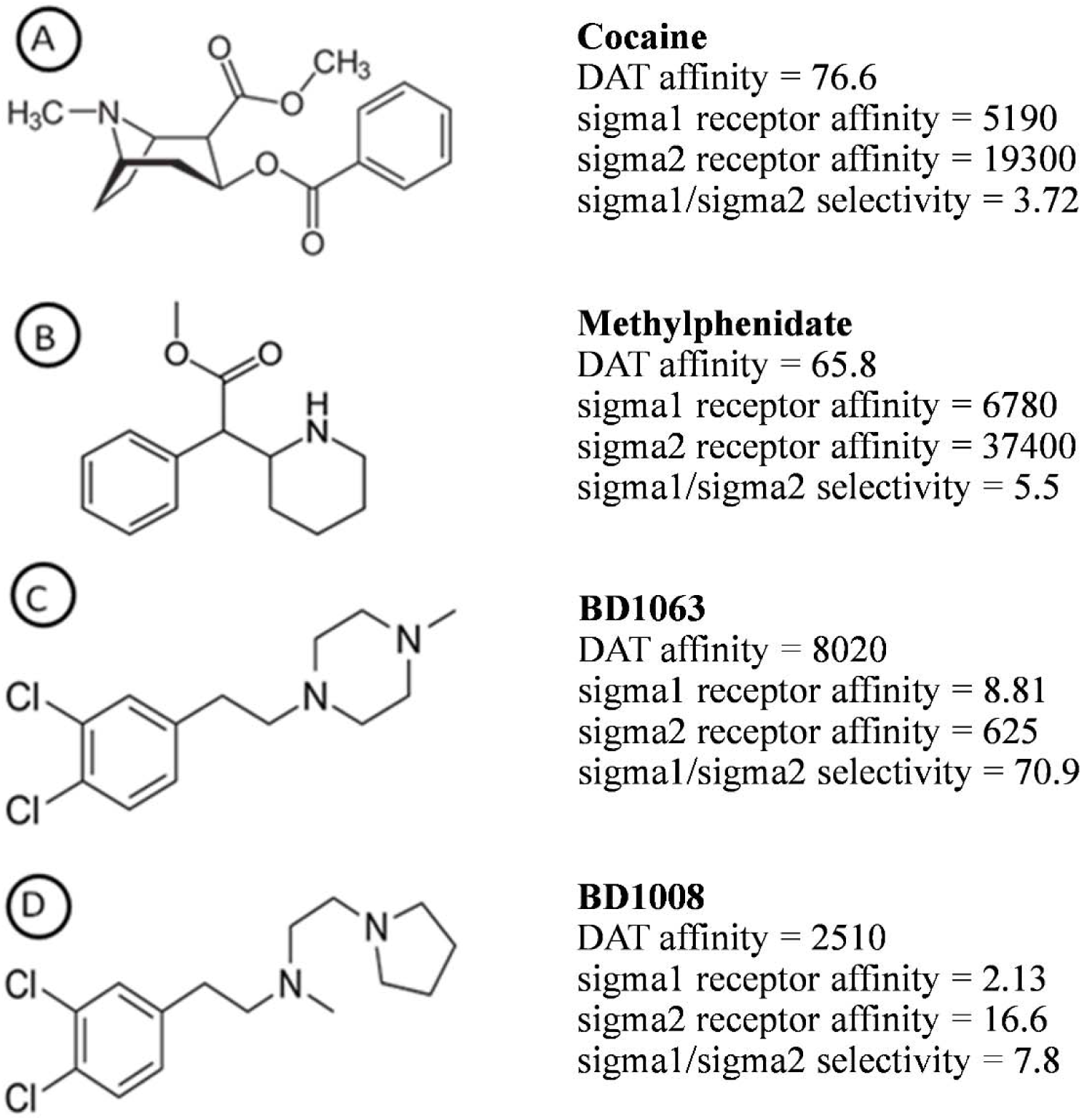
The drugs used in this study and their DAT and sigma1/sigma2 selectivity profiles. The DAT, sigma1 and sigma2 Ki were obtained from several references focused on rat brain tissue (Garcés-Ramrez et al., 2011; Hiranita et al., 2014; Kopajtic et al., 2010).

The BEST model, as the name implies, detects the ‘shift’ in the relationship between baseline and drug effects. This model involves a plot of baseline activity on the x-axis and a plot of drug activity on the y-axis, with the baseline-effect relationships fitted using linear (Figure 2A) or non-linear (Figure 2B) regression models. We can track if there is a shift in the baseline-effect relationship. A ‘shift’ would be confirmed if the difference between the curves is significant (P < 0.05). In principle, the BEST model can distinguish the effects of different drugs (Figure 2).

**Figure 2.**
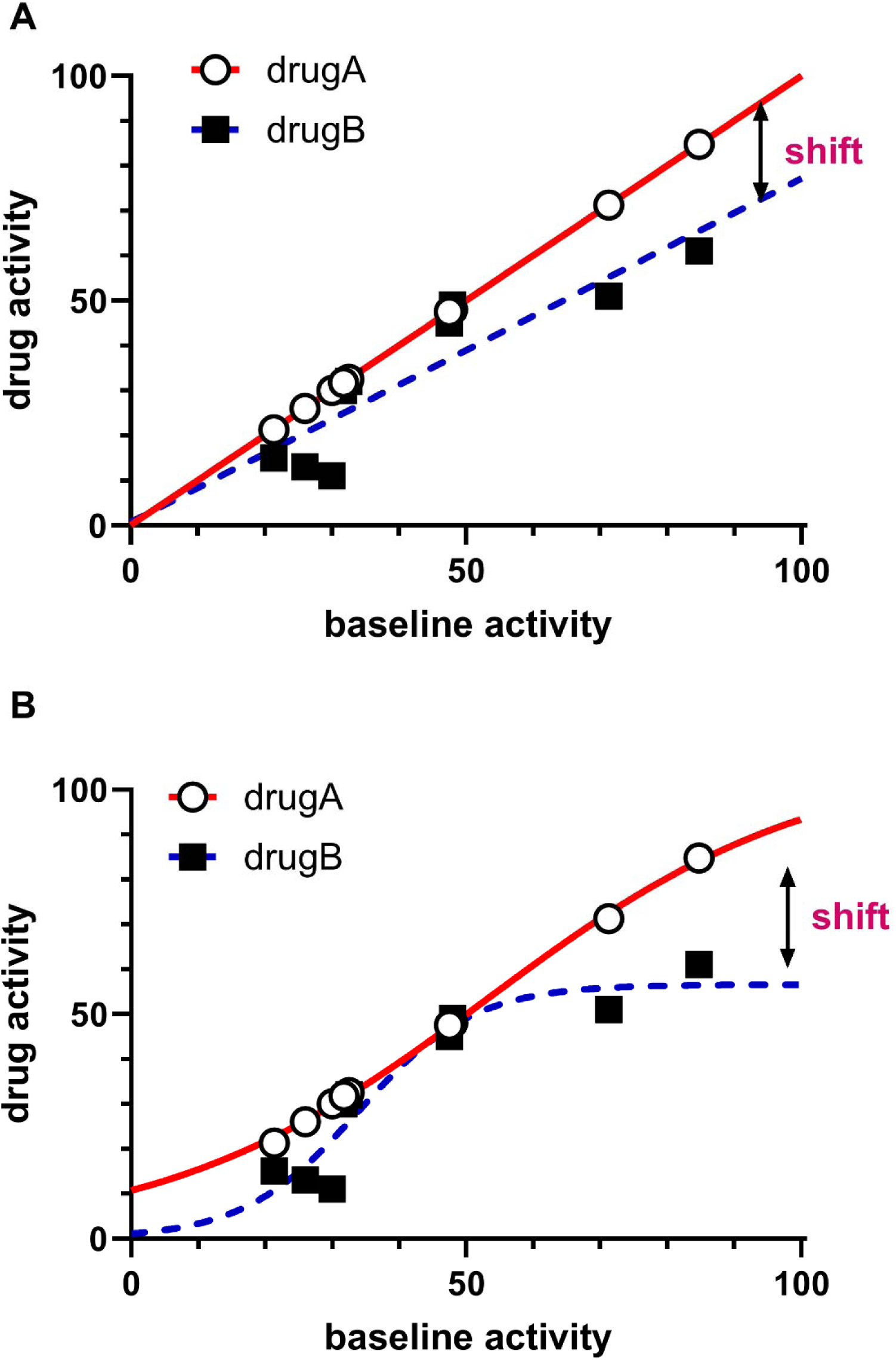
The baseline-effect-shift tracking (BEST) model. The following principles govern this model – 1) baseline activity is related to subsequent drug activity, 2) for two drugs that have similar baseline activity, the baseline-effect relationship may differentiate these two drugs in what we term a ‘shift’ which is the difference between the curves (see A-B). The same data is used in Fig A and B. The baseline-effect relationship is plotted with baseline activity on the x-axis and drug effect on the y-axis. The plot can be fitted with a linear (Fig A) or a non-linear curve (Fig B). The R^2^ values for drug A and drug B for the linear fit in Fig A were 1.0 and 0.81, respectively. The R^2^ values for drug A and drug B for the non-linear fit in Fig B were 1.0 and 0.90, respectively. It is important to use the best fit model – for example, the analysis of the data above revealed that the shift was not significant when we employ the linear plot in Fig A (F 1, 14 = 2.771, P = 0.1182), but for the same data set, the shift was significant is we employed the non-linear curve in Fig B (F 3, 12 = 10.16, P = 0.0013).

The goal of this study was to validate the BEST model by comparing it with the current model to determine which was more effective in distinguishing the effects of (MPD + BD1063) and (MPD + BD1008) on cocaine consumption under the constraints of price. To do this, we reanalyzed data presented previously, specifically the data with cocaine dose = 0.32 mg/kg/infusion wherein both of these drug combinations had no significant effects on cocaine consumption, and we included appropriate controls (20 mg sucrose/delivery) (Job and Katz, 2019). Our methods, results, discussion and conclusions are as below.

## Methods and Materials

For an overview of Methods, see (Job and Katz, 2019). All animal handling procedures and treatment schedules were approved by the National Institute on Drug Abuse Animal Care and Use Committee and followed the guidelines outlined in the National Institutes of Health (NIH) *Guide for the Care and Use of Laboratory Animals*.

### Drug combinations

The details are in shown in Table 1.

**Table 1.** Experimental design. For the experimental design, we used the same subjects (n = 9 for cocaine, n = 7 for sucrose) for all drug combinations and dose groups. Drug combinations included (MPD and BD1063) and (MPD and BD1008) each with different dose groups as shown above. The doses are in mg/kg and drugs were administered via the intraperitoneal route 5 min before behavioral economic analysis of demand curves using a within-session design as described in Methods section.

| Drug combination | (MPD + BD1063) | (MPD + BD1008) |
| --- | --- | --- |
| Dose group |  |  |
| 0 | MPD-0.0 + BD1063-0.0 | MPD-0.0 + BD1008-0.0 |
| 1 | MPD-1.0 + BD1063-0.0 | MPD-1.0 + BD1008-0.0 |
| 2 | MPD-1.0 + BD1063-3.2 | MPD-1.0 + BD1008-3.2 |
| 3 | MPD-1.0 + BD1063-10.0 | MPD-1.0 + BD1008-10.0 |

### Variables

Consumption at zero cost (Q_0_): We utilized behavioral economic analysis of demand curves to derive the variables corresponding to consumption at zero price (Q_0_) and the demand elasticity (alpha, α) (Figure 3A) using the exponential model (Hursh and Silberberg, 2008) as shown below:

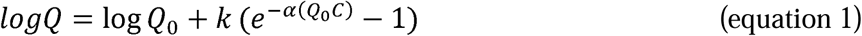

**Figure 3.**
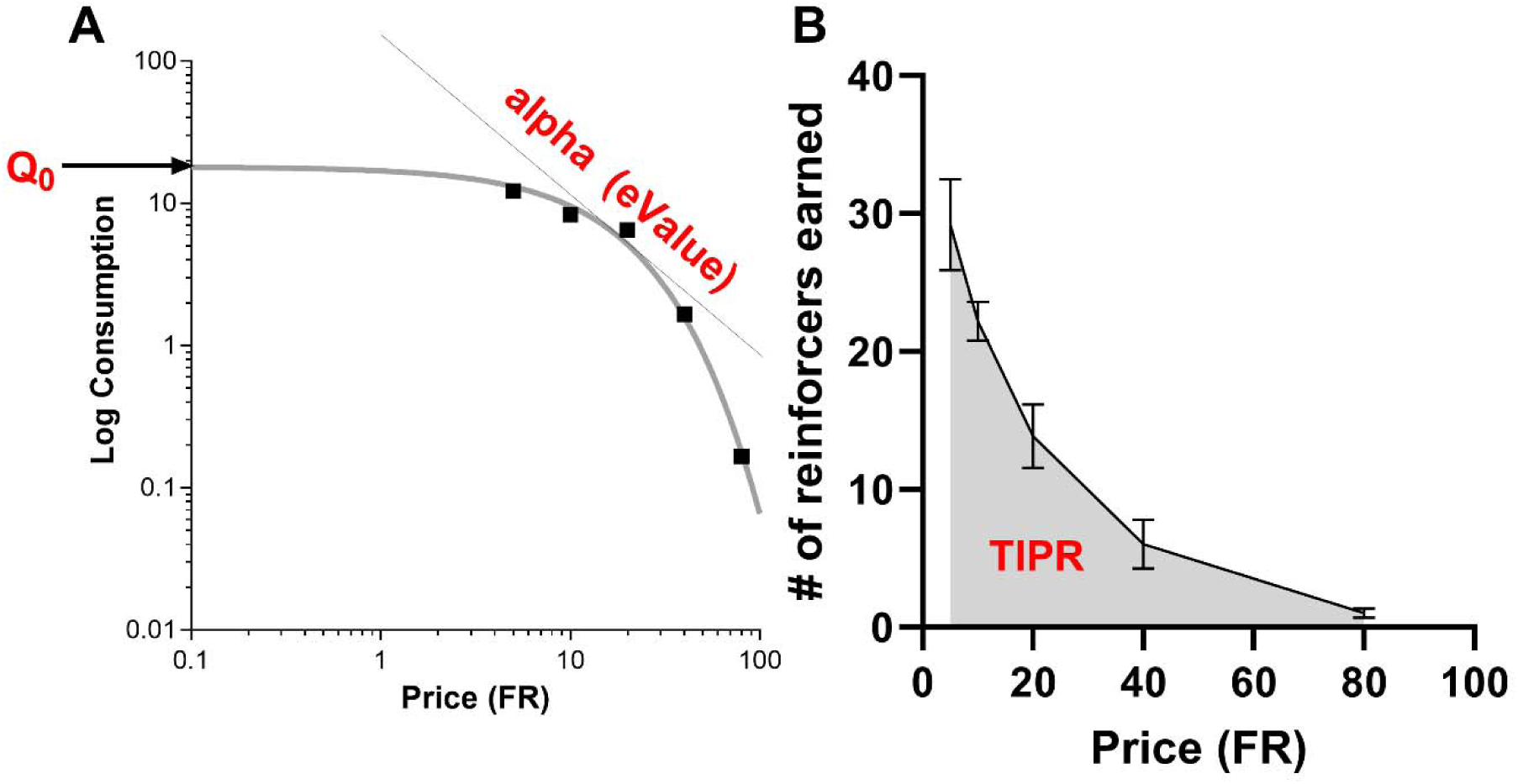
The variables. The variables used in this study include the behavioral economic derived parameters (see exponential model equation 1) corresponding to 1) the consumption at zero price (or Q_0_) which is the y-intercept of the demand curve in Fig A, and 2) the essential value (or eValue) which we derived, using equation 2, from the variable termed alpha or demand elasticity which represents how the curve bends towards the x-axis with increases in price. We also included the area under the curve of the price-response curve (Fig B). This is a sum of the number of reinforcers earned for the duration of all price increments for the completion of a within-session demand curve assessment.

Where Q is consumption, C is Price (FR requirements), Q_0_ is the consumption at minimal price (consumption when price = 0), α is a representation of the decline in consumption with increases in price, k is a scaling constant that is related to the consumption range. For this study we used k = 4.

Essential value (eValue): From the value of α derived from equation 1 above, we determined the essential value (eValue) using the following equation:

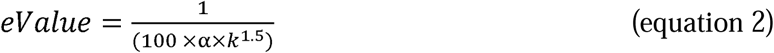

Where α and k are as described in equation 1. eValue is directly related to the motivation for the reinforcer.

Total intake under the price response curve (TIPR): We also calculated the area under the curve (AUC) of the price-response curve. The AUC is the sum of reinforcers earned under the constraints of price and we termed this total intake under the price response curve or TIPR (Figure 3B).

### BEST model

We plotted a graph of individual subject variables obtained from dose group 0 (control, see Table 1) on the x-axis and variables from dose groups 1-3 (see Table 1) on the y-axis for both drug combinations and employed regression analysis to fit these relationships. For these relationships, we employed linear (equation 3) and non-linear regression models (equation 4) for curve fit. The equation for linear fit is as follows:

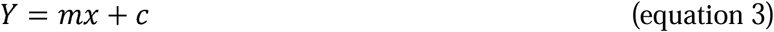

Where m = slope of the straight line, c is the y-intercept.

For non-linear curve fit we employed the sigmoidal function with the equation below:

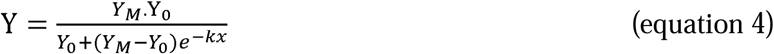

Where Y_M_ = maximum drug activity, Y_0_ = drug activity at baseline activity = 0, k = rate constant.

### Statistical analysis

We employed GraphPad Prism version 10 (GraphPad Software, San Diego, CA) and JMP version 18 (SAS Institute Inc., Cary, NC). For behavioral economic analysis of demand curves to obtain Q_0_ and alpha, we used the GraphPad template for the Exponential Model of Demand (equation 1) downloaded from the website: https://ibrinc.org/behavioral-economics-tools/. We calculated eValue using equation 2. We estimated TIPR as AUC, see above. Grubb’s test was used to determine if there were any significant outliers. Statistical significance was set at P < 0.05 for all analyses. For BEST model curves, we employed linear fit (equation 3) for Q_0_ and eValue and sigmoidal fit (equation 4) for TIPR. We compared the curves for the baseline-effect relationship for the different drug combinations. We employed normal mixtures clustering, where appropriate, to identify different groups of subjects based on baseline variable values. Two-way repeated measures ANOVA (factors are drug combination and dose group), One-way repeated measures ANOVA for each drug combination (factor = dose group) and linear regression analysis were employed to determine drug combination × dose group interaction, dose group effect and dose-dependency, respectively. We normalized our data by expressing as % change relative to baseline levels to allow more standardized comparisons. We used Dunnett’s test or Tukey’s post hoc tests, where applicable. Data are expressed as mean ± SEM.

## Results

### BEST model detects differences in the effects of the drug combinations

The baseline was the same as dose group 0. For (MPD + BD1063) versus (MPD + BD1008), there were no differences in the relationship between baseline Q_0_ and dose group 0 (Figure 4A). Furthermore, these curves were not different for the relationships between baseline Q_0_ and Q_0_ for dose group 1 (F 1, 14 = 1.900, P = 0.1897, Figure 4B), dose group 2 (F 1, 14 = 1.535, P = 0.2357, Figure 4C) and dose group 3 (F 1, 14 = 0.03233, P = 0.8599, Figure 4D).

**Figure 4.**
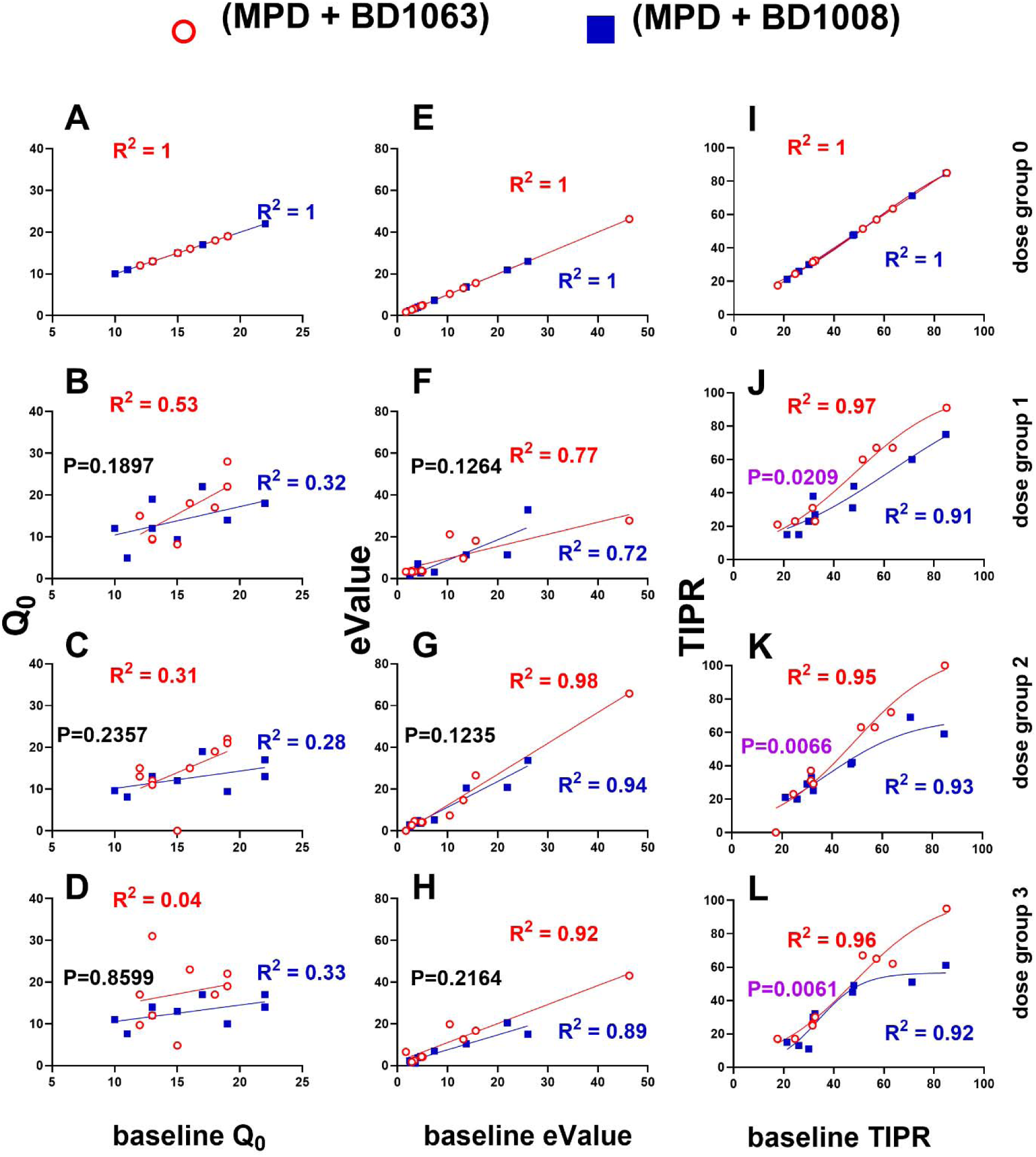
The BEST model distinguishes the effects of drug combinations (MPD and BD1063) and (MPD and BD1008) on total intake under the price response curve (TIPR) for cocaine consumption. The data points represent individual subjects with red opened circles representing (MPD + BD1063) and blue closed squares representing (MPD + BD1008) groups. The same subjects (n = 9) were used for all treatments. The left (Figure A-D), middle (Figure E-H) and right (Figure I-L) columns of graphs represent variables Q_0_, eValue and TIPR, respectively. The first, second, third and fourth rows represent drug dose group 0, 1, 2 and 3, respectively (see Table 1 for descriptions of dose groups). The x-axis represents baseline values for the appropriately labeled variables. The y-axis represents variable values for appropriately labeled dose groups. Linear regression was used to fit the curves for Figures A-H and sigmoidal fit was used to fit the curves for Figures I-L. Each graph shows the R^2^ values for curve fit using linear regression and P values for comparison between curves. A ‘shift’ is said to be detected when a comparison between the curves in a graph show P < 0.05. As shown above, there were no observed shifts for the relationships between baseline Q_0_ and baseline eValue and corresponding variables corresponding to dose groups (Figures A-H). However, there were shifts detected for the relationships between baseline TIPR and TIPR due to dose groups 1-3 (Figures J-L). In summary, BEST model detected a difference in the effects of (MPD + BD1063) and (MPD + BD1008), but only for TIPR, not for Q_0_ or eValue.

For eValue, (MPD + BD1063) versus (MPD + BD1008), there were no differences in the relationship between baseline and dose group 0 (Figure 4E). Additionally, there were no differences between these curves for the relationships between baseline eValue and dose group 1 eValue (F 1, 14 = 2.641, P = 0.1264, Figure 4F), dose group 2 eValue (F 1, 14 = 2.686, P = 0.1235, Figure 4G) and dose group 3 eValue (F 1, 14 = 1.676, P = 0.2164, Figure 4H).

For TIPR, we employed both linear and sigmoidal fit with details shown in Table 2. As shown from these, the sigmoidal fit resulted in better R^2^ values than the linear fit, and we employed this instead. For (MPD + BD1063) versus (MPD + BD1008), there were no differences in the relationship between baseline TIPR and dose group 0 (Figure 4I), as expected (F 3, 12 = 0.1538, P = 0.9252). However, these curves were different for baseline TIPR versus dose group 1 (Figure 4J), 2 (Figure 4K) and 3 (Figure 4L).

**Table 2.** A comparison between linear and sigmoidal fit. The table shows the R2 values for the curve fit for both linear and sigmoidal fit for the same data for dose groups 1-3 and also the P values for comparisons between the curves for the different drug combinations. Sigmoidal fit is more sensitive than linear fit in detecting differences between the drug combinations for the dose groups. Significance occurs when P < 0.05 (bolded).

|  | Linear fit | Sigmoidal fit |
| --- | --- | --- |
| Dose group 1 | Curve differences: (F 1, 14 = 3.410, P = 0.0860) | Curve differences: (F 3, 12 = 4.749, **P = 0.0209**, Figure 4J) |
|  | R<sup>2</sup> for (MPD + BD1063) = 0.96 | R<sup>2</sup> for (MPD + BD1063) = 0.97 |
|  | R<sup>2</sup> for (MPD + BD1008) = 0.92 | R<sup>2</sup> for (MPD + BD1008) = 0.91 |
| Dose group 2 | Curve differences: (F 1, 14 = 21.60, **P = 0.0004**) | Curve differences: (F 3, 12 = 6.690, **P = 0.0066**, Figure 4K) |
|  | R<sup>2</sup> for (MPD + BD1063) = 0.97 | R<sup>2</sup> for (MPD + BD1063) = 0.95 |
|  | R<sup>2</sup> for (MPD + BD1008) = 0.87 | R<sup>2</sup> for (MPD + BD1008) = 0.93 |
| Dose group 3 | Curve differences: (F 1, 14 = 7.463, **P = 0.0162**) | Curve differences: (F 3, 12 = 6.860, **P = 0.0061**, Figure 4L) |
|  | R<sup>2</sup> for (MPD + BD1063) = 0.95 | R<sup>2</sup> for (MPD + BD1063) = 0.96 |
|  | R<sup>2</sup> for (MPD + BD1008) = 0.81 | R<sup>2</sup> for (MPD + BD1008) = 0.92 |

### Normal mixtures clustering analysis detects two groups of subjects based on variable baselines

BEST model detected differences in the relationship between baseline variables and variables after drug treatment for at least one variable (TIPR) and the curve for (MPD + BD1008) for dose group 3 appears to represent a biphasic curve (Figure 4L). Because biphasic curves may imply more than one relationship between baseline variables and variables after drug treatment, we decided to explore for the possibility of more than one group in our sample. Because of the limitations of median split, we opted for a normal mixtures clustering analysis (Castaneda and Job, 2024).

Because we used the same set of animals twice, there were baseline variables for experiments with (MPD + BD1063) and for experiments with (MPD + BD1008). Normal mixtures clustering of the baseline values of all variables (Q_0_, eValue and TIPR) yielded two clusters (Figure 5A-B). Cluster1 (green) consisted of n = 5 subjects whereas cluster2 (red) consisted of n = 4 subjects. While there were no correlations between baseline Q_0_ for both experiments (Figure 5C), there were correlations between baseline TIPR and eValue (P < 0.05, Figure 5D-E).

**Figure 5.**
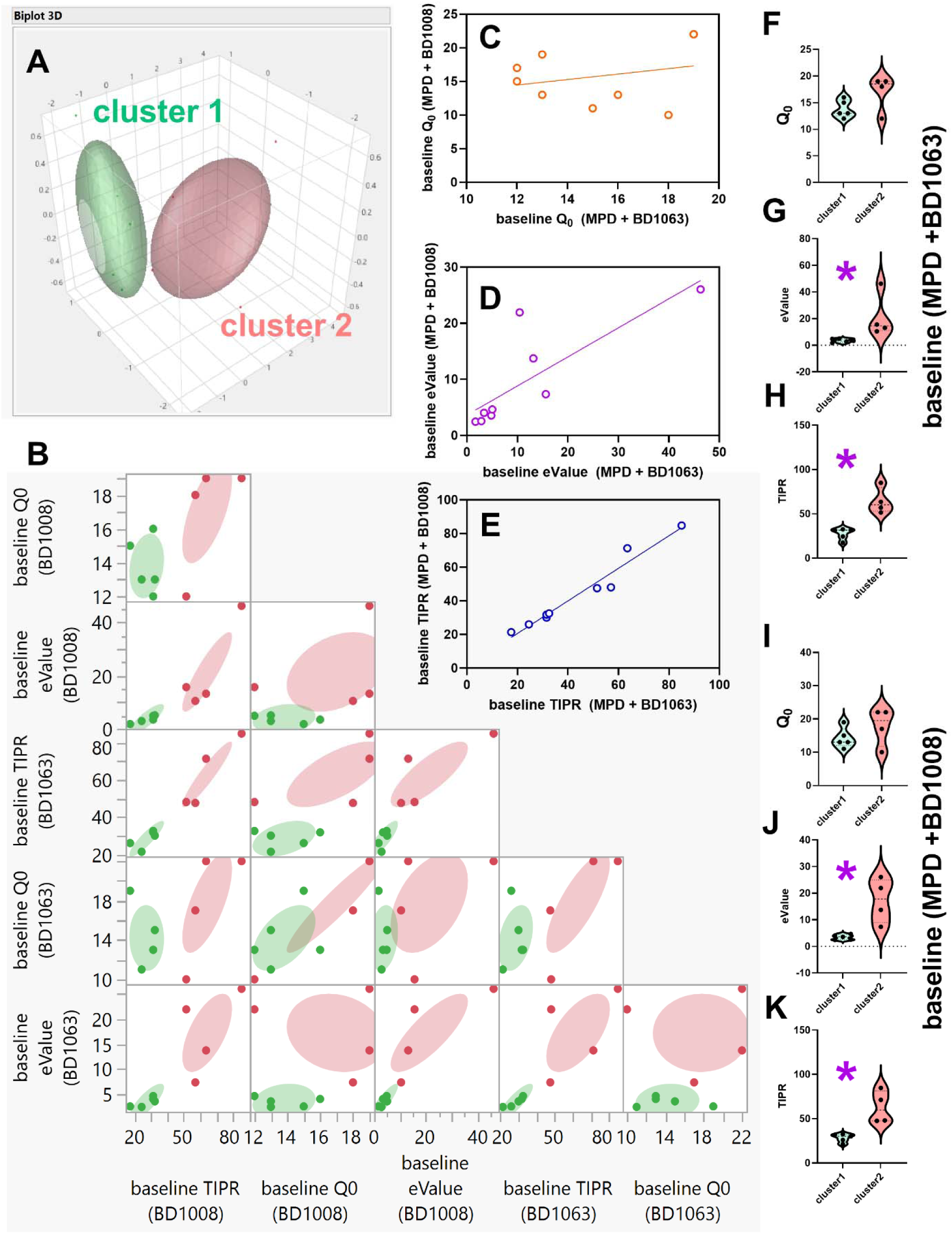
Normal mixtures clustering of all the values of baseline Q_0_, eValue and TIPR for experiments to determine the effects of (MPD + BD1063) and (MPD + BD1008) revealed two groups of subjects. For n = 9 subjects, the above procedure revealed two clusters (Figure A-B) which we named cluster 1 (n = 5, green) and cluster 2 (n = 4, red). Each data point represents an individual. For every individual we had 6 data points: baseline Q_0_, eValue and TIPR for each of the drug combination treatment experiments (MPD + BD1063) and (MPD + BD1008). The baseline values for Q0 for one drug combination experiment was not correlated with baseline values of Q_0_ for the other drug combination experiment (Figure C), but baseline eValue (Figure D) and TIPR (Figure E) were. These clusters identified were not distinct with regards to baseline Q_0_ (Figure F and I) (P > 0.05) but were distinct with regards to baseline eValue (Figure G and J) and baseline TIPR (Figure H and K). * show significant differences at P < 0.05.

The baseline Q_0_ for cluster1 and cluster2 for (MPD + BD1063) were 13.80 ± 0.73 and 17.0 ± 1.68, respectively. The baseline Q_0_ for cluster1 and cluster2, respectively, for (MPD + BD1008) were 14.20 ± 1.36 and 17.75 ± 2.84. Unpaired t-tests showed that cluster1 and cluster2 were not significantly different with regards to baseline Q_0_ for experiments for (MPD + BD1063) (P = 0.1013, Figure 5F) and experiments for (MPD + BD1008) (P = 0.2650, Figure 5I).

The baseline eValue for cluster1 and cluster2, respectively, for (MPD + BD1063) were 3.54 ± 0.62 and 21.37 ± 8.38. The baseline eValue for cluster1 and cluster2, respectively, for (MPD + BD1008) were 3.45 ± 0.42 and 17.27 ± 4.18. Unpaired t-tests showed that cluster1 and cluster2 were significantly different with regards to baseline eValue for experiments that included (MPD + BD1063) (P = 0.0466, Figure 5G) and for experiments that included (MPD + BD1008) (P = 0.0073, Figure 5J).

The baseline TIPR for cluster1 and cluster2, respectively, for (MPD + BD1063) were 27.50 ± 2.88 and 64.25 ± 7.34. The baseline TIPR for cluster1 and cluster2, respectively, for (MPD + BD1008) were 28.32 ± 2.08 and 62.88 ± 9.16. Unpaired t-tests showed that cluster1 and cluster2 were significantly different with regards to baseline TIPR for experiments involving (MPD + BD1063) (P = 0.0014, Figure 5H) and for experiments involving (MPD + BD1008) (P = 0.0044, Figure 5K).

In summary, the clustering identified two groups of subjects (Figure 5A-B) that were distinct with regards to baseline TIPR (Figure 5H and K) and eValue (Figure 5G and J, but similar with regards to baseline Q_0_ (Figure 5F and I).

### Current model versus BEST model

For distinctions in the effects of the drug combinations, BEST model (Figure 2) detected differences (Figure 4) particularly with regards to the variable TIPR (Figure 3). Follow up clustering of baseline values of all variables (Q_0_, eValue, TIPR) yielded two distinct clusters of subjects: cluster1 and cluster2 (Figure 5). Thus, the current model includes drug combinations and dose groups (see Table 1) whereas the BEST model includes drug combinations, dose groups and clusters. We compared the current model and the BEST model.

Q_0_: For the current model, Two-way repeated measures ANOVA did not reveal a drug combination × dose group interaction (F 3, 24 = 1.137, P = 0.3541, Figure 6A). Per drug combination, we also conducted One-way repeated measures ANOVA with factor = dose group (4 levels) to determine the effect of this independent variable on the dependent variables. For (MPD + BD1063), there was no dose group effect (F 1.930, 15.44 = 0.6848, P = 0.5139). For (MPD + BD1008), there was no effect of dose group (F 2.209, 17.67 = 3.090, P = 0.0665). Linear regression did not reveal any differences for (MPD + BD1063) versus (MPD + BD1008) for the relationships between dose group and % change (from baseline) of Q_0_ (F 1, 68 = 2.202, P = 0.1425, Figure 6B). Even after applying BEST model and separating into clusters, there was no drug combination × dose group interaction for cluster1 (F 3, 12 = 0.6441, P = 0.6013, Figure 6C) and cluster2 (F 3, 9 = 1.605, P = 0.2557, Figure 6E). When we focused on each drug combination, for cluster 1, the analysis revealed no effect of dose group on Q_0_ for (MPD + BD1063) (F 1.606, 6.426 = 0.9911, P = 0.4019) and for (MPD + BD1008) (F 1.942, 7.769 = 1.492, P = 0.2822). When we focused on each drug combination, for cluster 2, the analysis revealed no effect of dose group on Q_0_ for (MPD + BD1063) (F 1.488, 4.463 = 1.464, P = 0.3100) and for (MPD + BD1008) (F 1.685, 5.054 = 1.730, P = 0.2634). Furthermore, linear regression did not reveal any differences for (MPD + BD1063) versus (MPD + BD1008) for the relationships between dose group and % change in Q_0_ for cluster1 (F 1, 36 = 2.202, P = 0.9106, Figure 6D) and cluster2 (F 1, 28 = 2.389, P = 0.1334, Figure 6F).

**Figure 6.**
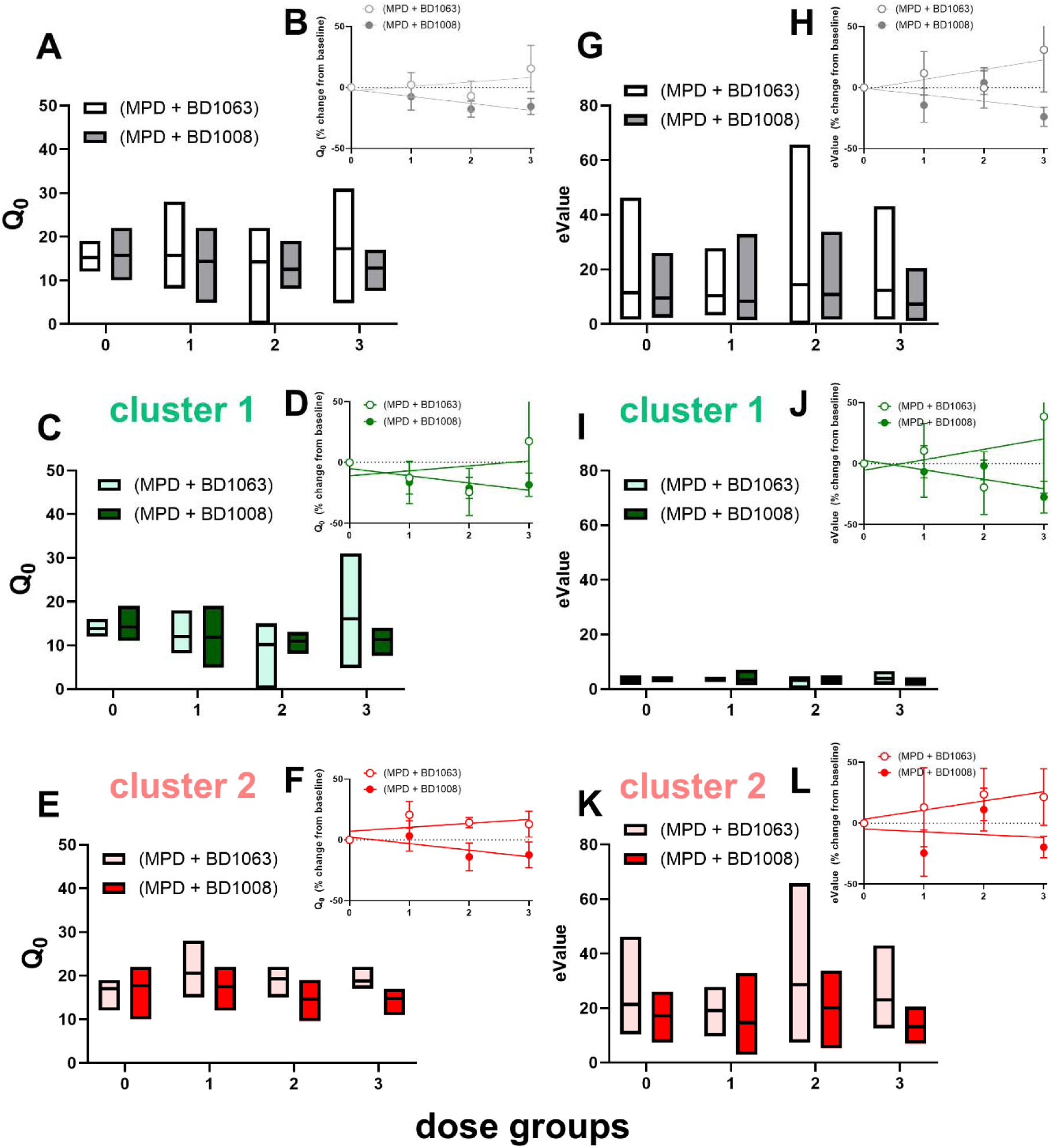
The current model and the BEST model do not distinguish the effects of drug combinations (MPD + BD1063) and (MPD + BD1008) and their dose groups on the variables Q_0_ and eValue for cocaine consumption. For all subjects (n = 9), Two-way repeated measures ANOVA with dependent variable Q_0_ and factors drug combination and dose groups yielded no drug combination × dose group interaction (Figure A). Linear regression did not reveal any differences for (MPD + BD1063) versus (MPD + BD1008) for the relationships between dose group and Q_0_ (expressed as % change from baseline Q_0_) (Figure B). After applying the BEST model, we identified two clusters – cluster1 (n = 5) and cluster 2 (n = 4). For cluster 1 (Figure C) and cluster 2 (Figure E), Two-way repeated measures ANOVA yielded no drug combination × dose group interaction. For cluster 1 (Figure D) and cluster 2 (Figure F), linear regression revealed no dose-dependent changes in Q_0_ (% change in Q_0_). The same lack of effects for Q_0_ were also observed for eValue (Figures G-L). In summary neither the current model nor the BEST model detected any differences in the effects of different dose groups of the drug combinations on Q_0_ and eValue.

eValue: For the current model, Two-way repeated measures ANOVA did not reveal a drug combination × dose group interaction (F 3, 24 = 0.3743, P = 0.7723, Figure 6G). For (MPD + BD1063), there was no dose group effect (F 1.172, 9.375 = 0.6995, P = 0.4464). For (MPD + BD1008), there was no effect of dose group (F 1.542, 12.33 = 1.541, P = 0.2493). Linear regression did not reveal any differences for (MPD + BD1063) versus (MPD + BD1008) for the relationships between dose group and % change (from baseline) for eValue (F 1, 68 = 1.734, P = 0.1923, Figure 6H). For BEST model-derived analysis, there was no drug combination × dose group interaction for cluster1 (F 3, 12 = 0.7477, P = 0.5442, Figure 6I) and cluster2 (F 3, 9 = 0.2421, P = 0.8648, Figure 6K). For (MPD + BD1063) cluster 1, there was no dose group effect (F 1.187, 4.747 = 0.1796, P = 0.7310). For (MPD + BD1008) cluster 1, there was no effect of dose group (F 1.579, 6.315 = 0.8771, P = 0.4357). For (MPD + BD1063) cluster 2, there was no dose group effect (F 1.105, 3.314 = 0.7635, P = 0.4551). For (MPD + BD1008) cluster 2, there was no effect of dose group (F 1.212, 3.635 = 1.258, P = 0.3469). Furthermore, linear regression did not reveal any differences for (MPD + BD1063) versus (MPD + BD1008) for the relationships between dose group and % change in eValue for cluster1 (F 1, 36 = 0.9851, P = 0.3276, Figure 6J) and cluster2 (F 1, 28 = 0.7354, P = 0.3984, Figure 6L).

TIPR: For the current method, a comparison of the drug combinations (2 levels) and dose groups (4 levels) for effects on TIPR using a Two-way repeated measures ANOVA did not reveal a drug combination × dose group interaction (F 3, 24 = 2.807, P = 0.0612, Figure 7A). For (MPD + BD1063) there was no dose group effect (F 2.322, 18.57 = 0.3857, P = 0.7151) whereas for (MPD + BD1008) there was a dose group effect (F 2.633, 21.07 = 4.043, P = 0.0240) with Dunnett’s test showing significant differences between dose group 0 versus dose group 1 (P = 0.0221) and between dose group 0 and dose group 3 (P = 0.0418). Linear regression analysis of the relationship between dose groups (x-axis) and average TIPR (% change from baseline) revealed a significant relationship for (MPD + BD1008) (F 1, 34 = 7.081, P = 0.0118) but not (MPD + BD1063) (F 1, 34 = 0.2342, P = 0.6316) and there were no differences between these slopes (F 1, 68 = 1.395, P = 0.2417, Figure 7B).

**Figure 7.**
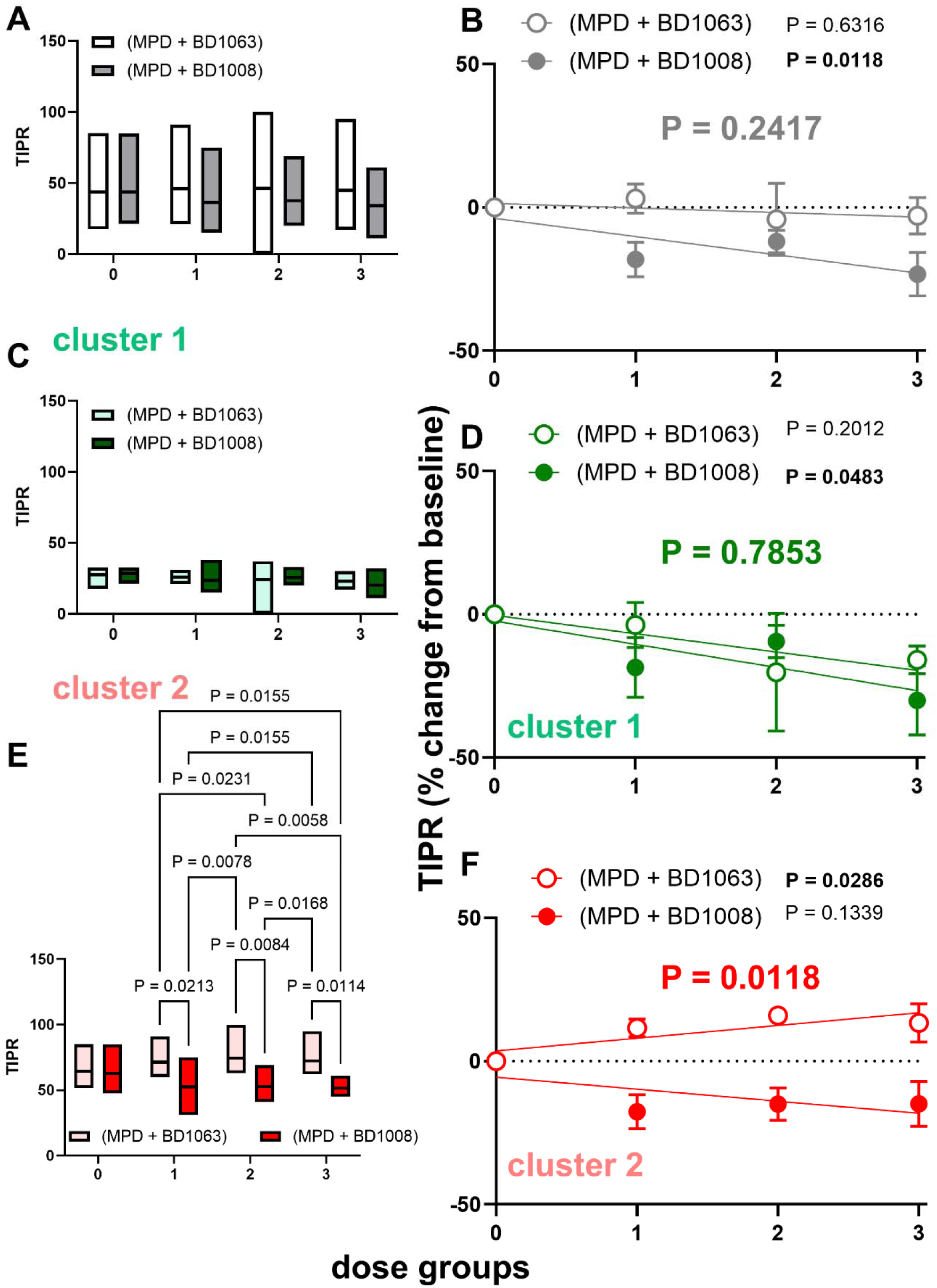
The BEST model, but not the current model, distinguishes the effects of drug combinations (MPD + BD1063) and (MPD + BD1008) and their dose groups on the variables TIPR in subjects with higher baseline consumption levels. For all subjects (n = 9), Two-way repeated measures ANOVA with dependent variable TIPR and factors drug combination and dose groups yielded no drug combination × dose group interaction (Figure A). Linear regression did not reveal any differences for (MPD + BD1063) versus (MPD + BD1008) for the relationships between dose group and TIPR expressed as % change from baseline TIPR (Figure B). However, for (MPD + BD1008) but not (MPD + BD1063) there was dose-dependency in its effects on % change in TIPR (Figure B). After applying the BEST model, we identified two clusters – cluster1 (n = 5, green) and cluster 2 (n = 4, red). For cluster 1 (Figure C), Two-way repeated measures ANOVA yielded no drug combination × dose group interaction and there were no differences in the slopes for dose-dependency assessments (Figure D). Like with the ungrouped sample in Figure B, for cluster 1, (MPD + BD1008) but not (MPD + BD1063) exerted a dose-dependent decrease in TIPR (expressed as % change in TIPR from baseline) (Figure D). For cluster 2, Two-way repeated measures ANOVA yielded a significant drug combination × dose group interaction (Figure E) and there were significant differences in the slopes for dose-dependency assessments (Figure F). For cluster 2, (MPD + BD1063) exerted a dose-dependent increase in TIPR (expressed as % change in TIPR from baseline) (Figure F) whereas (MPD + BD1008) exerted no significant effect (slope was not different from zero, Figure F). The P values following Tukey’s post hoc comparisons are written into the graph (Figure E). The P values for slope comparisons are written boldly into each of the graphs (Figure B, D and F). The P values for dose-dependency of each drug combination across dose groups are written next to the graph legends. In summary, combinations of (MPD + BD1008) appear to suppress cocaine TIPR more than (MPD + BD1063) whether subjects are ungrouped (Figure A-B) or separated into their respective clusters – cluster 1 (Figure C-D) and cluster 2 (Figure E-F). The drug effects, however, are significantly different for cluster 2 which corresponds to a group of subjects with higher baseline TIPR.

For BEST model, we conducted similar analysis but on each cluster. For cluster 1, with dependent variable = TIPR, a Two-way repeated measures ANOVA did not reveal a drug combination × dose group interaction (F 3, 12 = 0.4541, P = 0.7192, Figure 7C). For cluster 1, One-way repeated measures ANOVA revealed that there was no effect of dose group for (MPD + BD1063) (F 1.419, 5.678 = 0.6228, P = 0.5172) and (MPD + BD1008) (F 1.845, 7.379 = 2.605, P = 0.1410). Linear regression analysis of the relationship between dose groups (x-axis) and average TIPR (% change from baseline) revealed a significant relationship for (MPD + BD1008) (F 1, 18 = 4.487, P = 0.0483) but not (MPD + BD1063) (F 1, 18 = 1.760, P = 0.2012) and there were no differences between these slopes (F 1, 36 = 0.07532, P = 0.7853, Figure 7D). For cluster 2, a Two-way repeated measures ANOVA revealed a significant drug combination × dose group interaction (F 3, 9 = 5.235, P = 0.0230, Figure 7E) with Tukey’s post hoc tests showing significant differences between (MPD + BD1063) versus (MPD + BD1008) at every dose group (P < 0.05). One-way repeated measures ANOVA revealed an effect of dose group on TIPR for (MPD + BD1063) (F 1.845, 5.536 = 5.377, P = 0.05) with Dunnett’s test showing significant differences between dose group 0 versus dose group 1 (P = 0.0346) and between dose group 0 and dose group 2 (P = 0.0282). This was not observed for (MPD + BD1008) on cluster 2 as One-way repeated measures ANOVA did not show an effect of dose group (F 2.111, 6.332 = 1.866, P = 0.2310). Linear regression analysis of the relationship between dose groups (x-axis) and average TIPR (% change from baseline) revealed no significant relationship for (MPD + BD1008) (F 1, 14 = 2.532, P = 0.1339) but a significant relationship for (MPD + BD1063) (F 1, 14 = 5.954, P = 0.0286), and there were significant differences between these slopes (F 1, 28 = 7.264, P = 0.0118, Figure 7F).

BEST model did not detect any differences between the effects of the drug combinations on sucrose (Figure 8). There were no differences (P > 0.05) in the relationship between baseline variables and variables obtained for all the dose groups for the comparison between (MPD + BD1063) and (MPD + BD1008). Normal mixtures clustering of baseline variables revealed only one cluster (Figure 9A-B).

**Figure 8.**
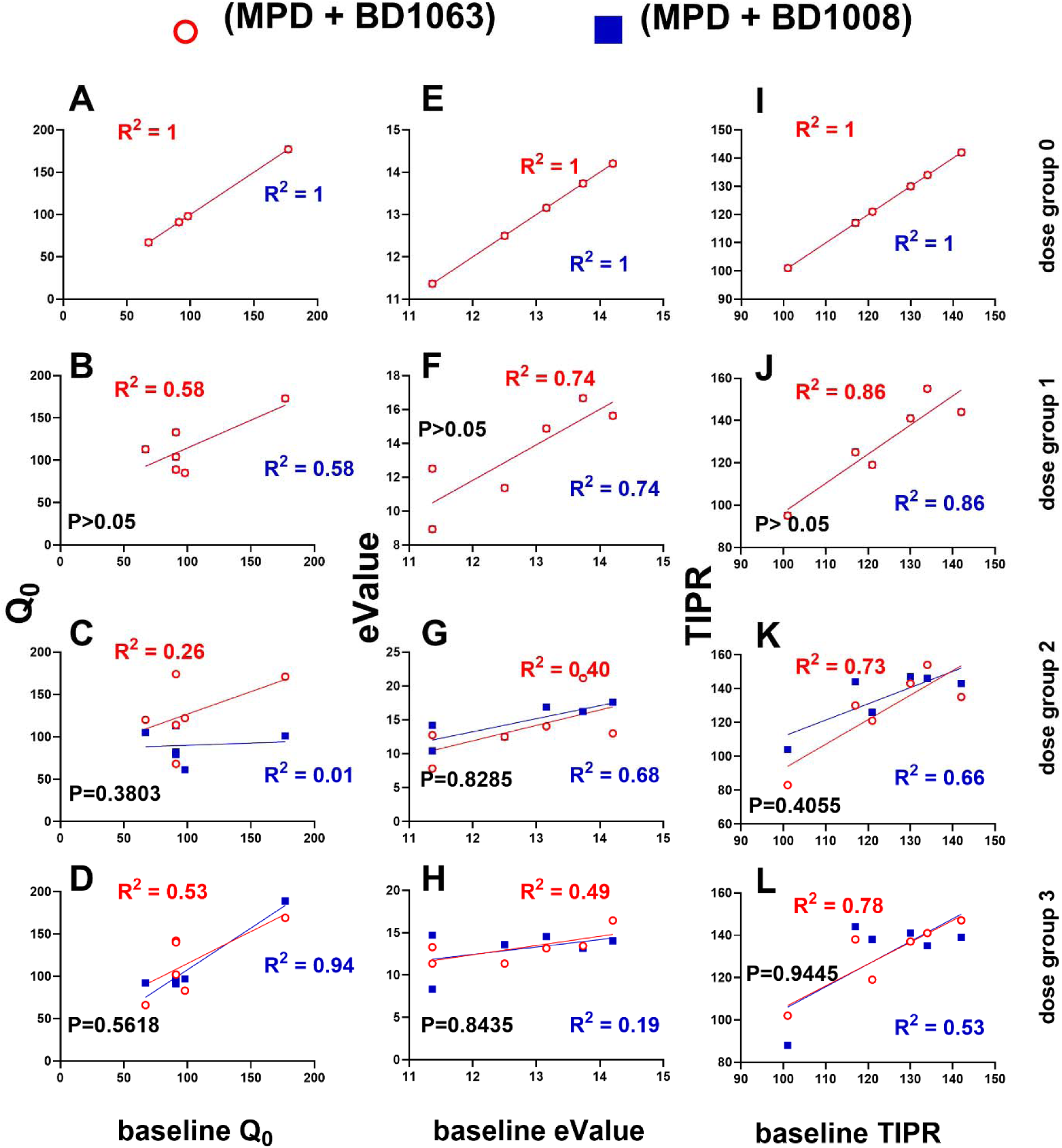
The BEST model did not distinguish the effects of drug combinations (MPD and BD1063) and (MPD and BD1008) on any variables for sucrose consumption. The data points represent individual subjects with red open circles representing (MPD + BD1063) and blue closed squares representing (MPD + BD1008) groups. The same subjects (n = 6) were used for all treatments. The left (Figure A-D), middle (Figure E-H) and right (Figure I-L) columns of graphs represent variables Q_0_, eValue and TIPR, respectively. The first, second, third and fourth rows represent drug dose group0, 1, 2 and 3, respectively (see Table 1 for descriptions of dose groups). The x-axis represents baseline values for the appropriately labeled variables. The y-axis represents variable values for appropriately labeled dose groups. Linear regression (see equation 3) was used to fit the curves. Each graph shows the R^2^ values for curve fit using linear regression and P values for comparison between curves. A ‘shift’ is said to be detected when a comparison between the curves in a graph show P < 0.05. As shown above, there were no observed shifts for any baseline-effect relationships for any dose groups. In summary, BEST model did not detect a difference in the effects of (MPD + BD1063) and (MPD + BD1008) for any variables associated with sucrose consumption.

**Figure 9.**
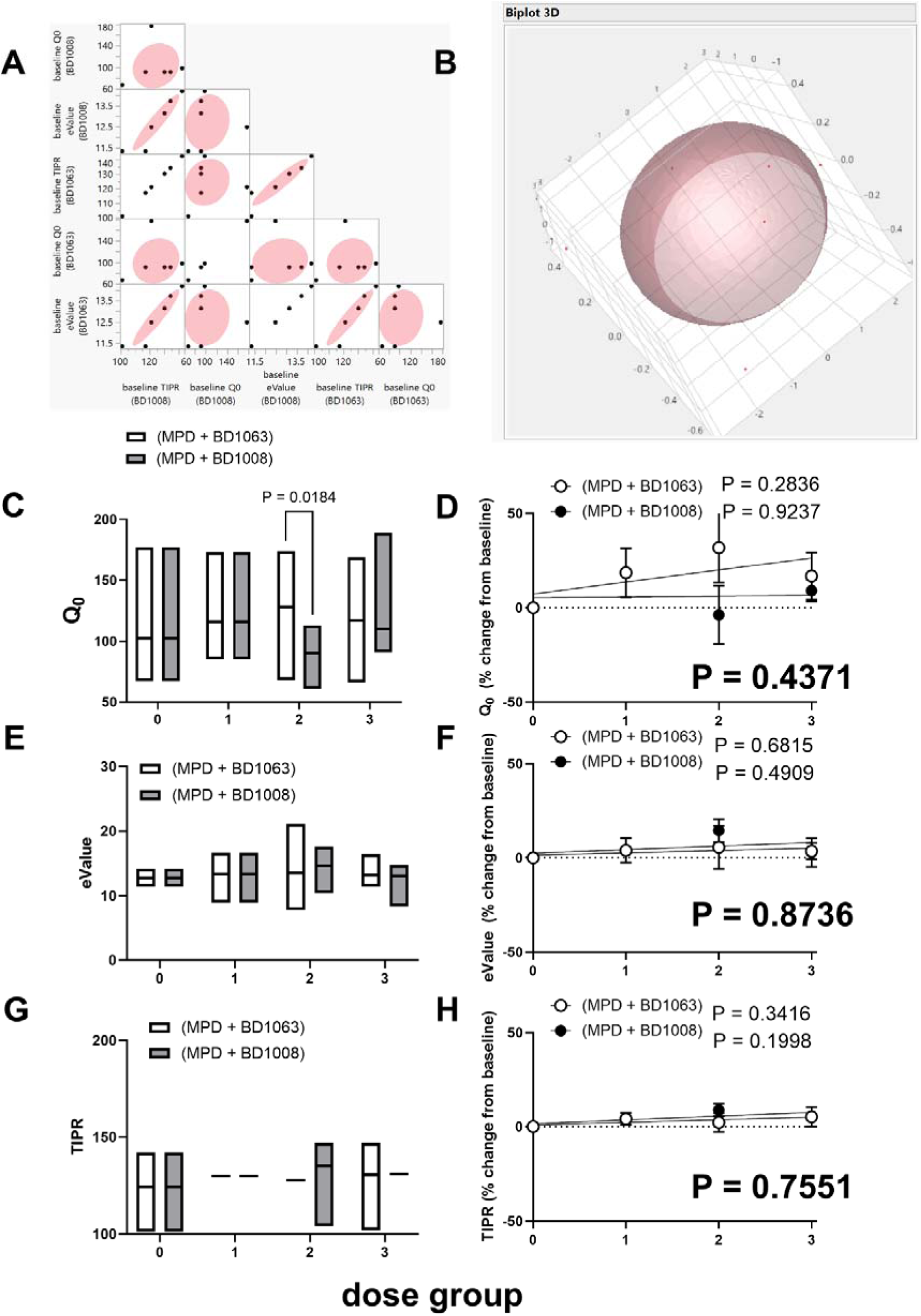
Current and BEST model do not distinguish the effects of drug combinations (MPD + BD1063) and (MPD + BD1008) and their dose groups on the variables eValue and TIPR for sucrose consumption: slight difference in their effects on Q_0_. Normal mixtures clustering revealed only one group of sucrose subjects (Figure A-B). For all subjects (n = 6), Two-way repeated measures ANOVA with dependent variable Q_0_ and factors drug combination and dose groups yielded a drug combination × dose group interaction (F 3, 15 = 3.708, P = 0.0354, Figure C) with Tukey’s post hoc test revealing significant differences for the drug dose 2. Linear regression did not reveal any differences for (MPD + BD1063) versus (MPD + BD1008) for the relationships between dose group and Q_0_ (expressed as % change from baseline Q_0_) (P = 0.4371, Figure D). For each drug combination, the slope was not different from zero (P > 0.05). For eValue (Figure E) and TIPR (Figure G), Two-way repeated measures ANOVA yielded no drug combination × dose group interaction (P > 0.05). Furthermore, linear regression did not reveal any differences for (MPD + BD1063) versus (MPD + BD1008) for the relationships between dose group and eValue (Figure F) and TIPR (Figure H) (both variables expressed as % change from baseline). The P values following Tukey’s post hoc comparisons are written into the graph (Figure C). The P values for slope comparisons are written boldly into each of the graphs (Figure D, F and H). The P values for dose-dependency of each drug combination across dose groups are written next to the graph legends (Figure D, F and H). In summary, there was no difference between these drug combinations with regards to dose-dependency (Figures D, F, and H) and no drug combination × dose group interaction for eValue (Figure E) and TIPR (Figure G), but a slight difference at a specific dose for Q_0_ (Figure C).

For Q_0_, a Two-way repeated measures ANOVA revealed a drug combination × dose group interaction (F 3, 15 = 3.708, P = 0.0354, Figure 9C) with Tukey’s post hoc test showing significant differences between the effects of (MPD + BD1063) and (MPD + BD1008) on Q_0_ only at dose group 2 (1 mg/kg of MPH and 3.2 mg/kg of drug). However, One-way repeated measures ANOVA did not show any effects of dose group for (MPD + BD1063) (F 2.445, 12.23 = 1.312, P = 0.3105) and (MPD + BD1008) (F 1.260, 6.300 = 1.887, P = 0.2220). Also, linear regression analysis did not show any significant differences between the regression lines (F 1, 44 = 0.6150, P = 0.4371, Figure 9D).

For eValue, a Two-way repeated measures ANOVA revealed a drug combination × dose group interaction (F 3, 15 = 0.4463, P = 0.7235, Figure 9E). One-way repeated measures ANOVA did not show any effects of dose group for (MPD + BD1063) (F 1.301, 6.503 = 0.1907, P = 0.7391) and (MPD + BD1008) (F 2.246, 11.23 = 2.335, P = 0.1385). Additionally, linear regression analysis did not show any significant differences between the regression lines (F 1, 44 = 0.02559, P = 0.8736, Figure 9F).

For TIPR, a Two-way repeated measures ANOVA did not reveal a drug combination × dose group interaction (F 3, 15 = 1.248, P = 0.3274, Figure 9G); One-way repeated measures ANOVA did not detect a dose group effect for (MPD + BD1063) (F 1.691, 8.455 = 0.9410, P = 0.4119) and (MPD + BD1008) (F 1.992, 9.960 = 1.736, P = 0.2255); linear regression analysis did not show any significant differences between the regression lines (F 1, 44 = 0.09849, P = 0.7551, Figure 9H).

The summary of the results is shown in Table 3.

**Table 3.** Summary of results (current versus BEST model comparisons) Under conditions where the drug combinations had no significant effects on cocaine consumption, BEST model was able to determine that there was a difference in the effect profiles of these drug combinations with regards to different types of responses in the sample. This speaks to the sensitivity of the BEST model over the current model, but also to the sensitivity of the variable TIPR over Q_0_ and eValue when it comes to distinguishing the effects on cocaine consumption of these closely related drug combinations. Cluster 2 have higher baseline cocaine consumption than cluster 1. NA means not applicable

| Current model | BEST model |
| --- | --- |
| NA | Detects differences in the relationship(s) between baseline variables and dose group effect on variables for cocaine self-administration under the constraints of price increases (Figure 4) |
| NA | Detects biphasic relationship later confirmed via normal mixtures clustering analysis as two types of behavioral responses in the sample linked to differences in baseline variables (Figure 5) |
| Detects no effect of drug combination 1 and drug combination 2 including different dose groups on the variables $Q_0$ and eValue (Figure 6), corroborated by previous study (Job and Katz, 2019) | Detects no effect of drug combination 1 and drug combination 2 including different dose groups on the identified groups for the variables $Q_0$ and eValue (Figure 6) |
| Detects no differences between the effects of drug combination 1 and drug combination 2 including different dose groups on the variable TIPR (Figure 7) | Detects differences between the effects of drug combination 1 and drug combination 2 including different dose groups on cluster 2 for the variable TIPR (Figure 7E-F) |
| NA | Detects no differences in the relationship(s) between baseline variables and dose group effect on variables for sucrose delivery (Figure 8-9) |

## Discussion

The goal of this study was to compare the current model to our new BEST model (Figure 2) to determine which was more effective in distinguishing the effects of combinations of (MPD + BD1063) and (MPD + BD1008) on cocaine consumption. Our important findings are two-fold. We show that 1) employing the new BEST model relative to the current model represents a more sensitive approach for distinguishing drug effects, and 2) the TIPR variable relative to Q_0_ and eValue represents a more sensitive variable, at least with respect to this study, for the purposes of distinguishing the effects of similar drugs on cocaine consumption.

The effects of these drug combinations are baseline-dependent. With reference to cocaine consumption under the constraints of price, and at the cocaine 0.32 mg/kg dose, drug combinations that include BD1063 can exert no effect and/or increase cocaine consumption, whereas the combinations containing BD1008 can decrease and/or have no effects – with these effects all depending on the baseline cocaine consumption level of the subject. This baseline-dependency of drug effects on psychostimulant consumption is buttressed by a recent report in which we showed that the effect of chemogenetically-mediated inhibition of dorsal striatum dopamine D1 receptor-related activity on methamphetamine (METH) intake was dependent on baseline METH experience in female Long Evans rats (Madhuranthakam and Job, 2024).

Similarly, subjects with different baseline experiences with METH (subjects taking METH under restricted/short access conditions versus subjects taking METH under extended access conditions) showed differential sensitivities to the suppressing effects of drugs that modulate the dopamine system on METH self-administration (Orio et al., 2010; Tunstall et al., 2018; Wee et al., 2007).

Because we identified distinct clusters, it is tempting to try to explain these clusters as different types of drug users. What is clear is that they represent differential responses of cocaine self-administration to the same drug combinations, but this was related to differential baseline cocaine consumption behavior. It is not clear that they are different types of drug users. Moreover, these clusters had the same Q_0_, and overall do not meet the requisite criteria we proposed in a previous study to qualify as different drug user types (Castaneda and Job, 2024).

Our results suggest that, combined with MPD, the non-selective sigma receptor antagonist (BD1008) was more effective at suppressing cocaine consumption than the selective sigma1 receptor antagonist (BD1063) in a group of subjects that had higher baseline cocaine consumption behavior. Our results are similar to that obtained from a previous study that showed that a drug combination of a dopamine transporter inhibitor (WIN 35,428, 0.1 mg/kg and 0.3 mg/kg) and BD1063 at similar doses to that which we used in this study (3.2, 10 mg/kg), did not attenuate cocaine (0.32 mg/kg/injection) self-administration while the same drug combination with BD1008 did (Hiranita et al., 2011).

The differences in BD1008 versus BD1063 effects on cocaine consumption may have something to do with dopamine release. For example, BD 1008 suppresses dopamine release in the mesolimbic dopamine system while BD1063 does not (Garcés-Ramrez et al., 2011). Also, because drug combinations with BD1008 have more affinity at sigma2 than drug combinations with BD1063, the differences in their effects on cocaine consumption in subjects with higher baseline cocaine consumption levels (Table 2) may be because cocaine consumption at this higher level involves sigma2 > sigma1 receptor input. Indeed, there is evidence that under conditions of repeated cocaine exposure, there is a transition from predominantly sigma1 receptor-related mechanisms for dopamine activity regulation to sigma2 receptor-related mechanisms of dopamine activity regulation in the dorsal striatum (Aguinaga et al., 2018).

A limitation of this work is that we did not examine sex differences in the effects of these drug combinations on cocaine consumption. With increasing interests in understanding sex as a biological variable (SABV), it will be imperative to conduct studies in both male and female subjects in future studies. Another limitation is that we did not explore other doses of MPD. We agree that other doses should be explored. That said, we show that for drug combination, including doses, that have not shown effectiveness in suppressing cocaine consumption, BEST model determined that this lack of effect was due to different effects on groups of subjects with different baseline cocaine consumption behavior.

BEST model (with TIPR analysis) is more sensitive than the current models in differentiating drug effects on cocaine consumption. The major implication of our results is that we can separate the effects of closely related drugs pharmacologically using our new BEST model, a finding that may be important to advance this field.

## Acknowledgement

The authors wish to acknowledge Dr. Jonathan L. Katz in whose laboratory MOJ conducted the behavioral experiments. All authors contributed to data analysis and to the writing of the manuscript. MOJ designed and conducted the behavioral experiments and statistical analysis. This work was funded by the Department of Health and Human Services/National Institutes of Health/National Institute on Drug Abuse/Intramural Research Program, Baltimore, MD, USA grant DA000547. This work was also supported by the Francis Lax Fund for Faculty Development at Rowan University. This work was also supported by startup funds from Rowan University, Camden, New Jersey.

## Disclosures

Reshma Paul has no conflicts of interest to declare. Roshni M Gandhi has no conflicts of interest to declare. Dr. Martin O Job has no conflicts of interest to declare.

## Notes

### Competing Interest Statement

The authors have declared no competing interest.

